# Classical conditioning in oddball paradigm: A comparison between aversive and name conditioning

**DOI:** 10.1101/286492

**Authors:** Yuri G. Pavlov, Boris Kotchoubey

**Author notes:** Corresponding author: Yuri G. Pavlov, Address: Institute of Medical Psychology and Behavioral Neurobiology, Silcherstr. 5, 72076, Tübingen, Germany.

## Abstract

The nature of cortical plasticity in learning is one of the most intriguing questions of the modern cognitive neuroscience. Classical conditioning (as a typical case of associative learning) and electroencephalography together provide a good framework for expanding our knowledge about fast learning-related cortical changes. In our experiment we employed a novel paradigm in which classical conditioning was combined with passive oddball. Nineteen subjects participated in the first experiment (aversive conditioning with painful shock as unconditioned stimulus, US and neutral tones as CS), and 22 subjects in the second experiment (with a subject’s own name as US). We used event-related potentials (ERP), time-frequency and connectivity analyses to explore the CS-US interaction. We found a learning-induced increment of P3a in the first experiment and the late positive potential (LPP) in both experiments. These effects may be related to increased attentional and emotional significance of conditioned stimuli. We showed that the LPP and P3a effects, earlier found only in visual paradigms, generalize to the auditory sensory system. We also observed suppression of the low beta activity to CS+ in aversive conditioning over the contralateral to expected electrical shocks hemisphere, presumably indicating preparation of the somatosensory system to the expected nociceptive US. No evidence of increased connectivity between somatosensory (representing painful US) and auditory (CS) cortex was found.

## Introduction

Aversive conditioning^1^ has been an object of numerous psychophysiological studies, many of which used EEG and event-related potentials (ERPs: reviews in Christoffersen & Schachtman, 2016; Miskovic & Keil, 2012). Most studies employed visual conditioned stimuli (CS), typically paired with nociceptive unconditioned stimuli (US) (e.g. Hermann, Ziegler, Birbaumer, & Flor, 2000; Pizzagalli et al., 2003; Wong et al., 2004). This combination has several advantages, for example well-manifested pattern of EEG/ERP responses and the fast rate of conditioning (e.g., Olofsson, Nordin, Sequeira, & Polich, 2008). There are also some disadvantages, however. We live in a multisensory environment but studies of other sensory modalities in this field are still uncommon. Only two ERP studies applied pure auditory CS with nociceptive US (Kluge et al., 2011; Waschulewski-Floruss, Miltner, Brody, & Braun, 1994). The effects of conditioning on P3 (Baas, Kenemans, Böcker, & Verbaten, 2002; Begleiter & Platz, 1969; Franken, Huijding, Nijs, & van Strien, 2011; Wong et al., 2004) and the late positive potential (LPP) were demonstrated in responses to visual CS (Bacigalupo & Luck, 2018; Hermann et al., 2000; Wong et al., 2004) but not to auditory CS.

Furthermore, complex visual stimuli can hardly be useful in low-responsive populations such as patients with severe brain damage, mentally ill, or small children. These individuals’ difficulties in visual perception are caused by their impaired gaze control or the inability to follow instruction to fix their gaze. To the contrary, hearing disorders are strongly limited to patients with selective lesions of auditory pathways, which is rather rare. Therefore, auditory paradigms are successfully employed in many groups of severely brain-damaged patients (for review of achievements see Kotchoubey, 2015; for future perspectives see Kotchoubey, Pavlov, & Kleber, 2015). Such paradigms can deliver a reliable measure for studying associative learning without any instruction in patients having no overt behaviour altogether.

Perception is facilitated when stimuli are predictable (Friston, 2005; Grossberg, 1982). A violation of the predictions engages additional neural resources to adjust the predictive model (Friston, 2005, 2010). An example of this process is the oddball paradigm where the appearance of a rare deviant in a sequence of frequent standard stimuli elicits a mismatch negativity (MMN) and P300 responses. P300 in passive oddball may reflect involuntary attention capture and conscious deviance detection (Schröger, 1997). We assumed that additional attentional resources engaged because of prediction errors and reflected in P300 can reinforce CS-US associations. That is, a deviant CS in the oddball conditioning paradigm sound would attract attention, thereby increasing the chance of the following US to get into the focus of attention.

Classical conditioning can also be seen as a predictive process (Anokhin, 1973). After multiple CS-US pairings the brain formulates certain expectations regarding CS. A violation of the expectations (in aversive conditioning, the omission of the noxious event) generates a prediction error signal. Already Durup & Fessard (1935) observed conditioned alpha suppression to the sound of a camera shutter in anticipation of the camera flash. More recently conditioned alpha suppression was demonstrated by Babiloni (2003) and Harris (2005), while other studies also showed suppressed EEG activity in the beta band (van Ede, de Lange, Jensen, & Maris, 2011; van Ede, Jensen, & Maris, 2010; van Ede, Szebényi, & Maris, 2014). On the other hand, stimulus omission can elicit suppression of the alpha (Andersen & Lundqvist, 2019) and beta activity (Moses, Bardouille, Brown, Ross, & McIntosh, 2010). These results indicate that an analysis of EEG oscillatory activity (in the alpha, beta) in associative learning may substantially complement and extend the knowledge obtained using ERPs.

To summarize, in an oddball conditioning paradigm a violation of the first order prediction (the oddball effect) may help to attract additional resources to process CS. This, in turn, activates the second order prediction of the following US. Taking advantage of the multisensory nature of the task we will be able to separate direct EEG/ERP responses to auditory CS, preparatory activity reflecting expectation of somatosensory aversive US, and violation of the prediction when US is omitted.

According to the ideas of Hebb (1949), associative learning should be related to formation of novel neuronal assemblies. This process can manifest itself in EEG phase synchronization (Fell & Axmacher, 2011). Specifically, cross-modal conditioning paradigms are best appropriate to disentangle phase synchronization related to perception from genuine formation of connections between brain areas. Thus increased coherence in the gamma band was found in visual-somatosensory aversive conditioning (Klein, Sauer, Jedynak, & Skrandies, 2006; Miltner, Braun, Arnold, Witte, & Taub, 1999). We used auditory-somatosensory cross-modal conditioning to see whether we can reproduce this effect in a different set of modalities.

A potential methodological problem of using ERPs to study conditioning is that ERPs need many trials to get a good signal-noise ratio, but conditioned response (CR) quickly extinguishes without reinforcement (Lonsdorf et al., 2017; Sperl, Panitz, Hermann, & Mueller, 2016). To study CR in reinforced trials, CS-US intervals must be long enough to prevent an overlap of responses to CS and US. Another solution is a partial reinforcement design. Skin conductance experiments (Bouton, Woods, & Pineño, 2004; Culver, Stevens, Fanselow, & Craske, 2018) showed that suspending of reinforcement leads to a rapid decrease of the CR amplitude, but that partial reinforcement with only 10 to 20% reinforced trials is sufficient to maintain the CR at the level attained during acquisition. Taking into account the tendency to fast attenuation of skin conductance responses (Bacigalupo & Luck, 2018) we expected that a similar technique would yield at least the same (or even better) results in ERP.

The first aim of the study was largely practical. We looked for a simple learning paradigm that might be reliably applied in various groups including low-responsive individuals whose attention, visual perception and overt behaviour are severely impaired. Aversive conditioning is a paradigm with highly reliable effects but these effects are mostly attained using nociceptive US that have both ethical and methodological problems. As regards the former, low-responsive individuals cannot give an informed consent. The consent given by their legal representatives permits to avoid legal issues, but from the ethical point of view it cannot fully replace the subject’s own statement. As regards the latter, there are considerable individual differences in pain sensitivity. Therefore, in normally responsive individuals nociceptive stimuli are usually adjusted to the individual sensation and pain thresholds, but this complex procedure requires a high level of cognitive functioning, making it impossible in individuals with restricted abilities.

Several authors suggested replacement of the strong aversive stimuli in sensible populations by a different kind of potentially highly significant stimuli, subjects’ own names (SON) (e.g., (Fischer, Dailler, & Morlet, 2008; Kotchoubey, Lang, Herb, Maurer, & Birbaumer, 2004; Perrin et al., 2006; Wang et al., 2015). The results of the use of SON as a meaningful stimulus are, however, not consistent. In our preliminary study (Kotchoubey & Pavlov, 2017) we showed a significant conditioning effect on ERPs using SON as reinforcement at the group level, but not as the individual level. In the present study we intended to directly compare strong aversive conditioning with conditioning based on SON reinforcement.

The second aim of the study was to assess the generalizability of ERP phenomena earlier demonstrated only in visual paradigms. We expected an increase of ERP components P3 and the LPP as signs of attention and emotional processing. This increase would be more pronounced in aversive conditioning than in SON conditioning.

The third aim was to explore the effects of expectancy violation on EEG oscillatory activity in the auditory-somatosensory cross-modal conditioning paradigm. The SON experiment in this respect would serve as a control. Because the nociceptive US is lateralized but the SON is not, we expected a lateralized activity in response to the auditory CS in the aversive conditioning experiment but not in the SON experiment.

The fourth aim was to approach mechanisms of multisensory integration. Thus we performed a connectivity analysis and hypothesised that aversive conditioning would enhance functional connections between the somatosensory and auditory cortex.

Finally, because the LPP effect is usually attributed to emotional factors, we expected that this effect would correlate with personality traits Anxiety and Neuroticism in Aversive conditioning but not in Name conditioning. However, taking into account a relatively small sample size in our study, we regard the results of the correlational analysis as preliminary; the details are presented in Supplementary materials.

## Methods

### Participants

Twenty-three healthy subjects participated in the Aversive conditioning experiment. One participant was excluded from the analyses due to excessive movement artefacts and three due to technical problems. The final sample included 19 participants (12 females; mean age = 24.63, SD = 2.29). 22 healthy individuals participated in the Name conditioning experiment (13 females, mean age 25.70, SD = 2.26). 17 participants took part in both experiments (11 females; mean age = 24.88, SD = 2.28).

None of the participants had had any disease of the nervous system or hearing disorders in the past, or reported use of any drugs during the last week before the experiment. Participants were seated in a comfortable chair and asked to close their eyes and to listen attentively to the stimuli. Informed consent was obtained from each participant. The study was approved by the Ethical Committee of the University of Tübingen.

### Stimuli and conditioning procedure

#### Aversive conditioning

Before the experiment we conducted a setting threshold procedure to adjust the amplitude of the electrical shock to an individual pain threshold. A single 50-μs electrical shock generated by Medicom MTD electrical stimulator was delivered to the left wrist. The stimulation was initially set at 1 mA and the intensity was gradually increased with 1 mA steps until the participant indicated that he or she sensed the stimulus. This point was regarded as a first sensory threshold. We continued increasing the intensity until the participant reported that at this level the electrical shock can be considered painful (“the slightest pain possible”). After this point (i.e., the first pain threshold), the level 80% above this threshold was reached in 5 linearly distributed steps. After the shock of 1.8 pain threshold we asked participants to assess the current stimulation level as bearable or too high. All participants reported the current level as moderately painful but not too strong. The procedure then was repeated in the opposite direction, decreasing the stimulation from the level of 1.8 pain threshold to the level at which the stimulus was not experienced as pain anymore (i.e., the second pain threshold), and further decreasing it to the level at which the participant ceased to experience the stimulus altogether (i.e., the second sensory threshold). The final values of the sensory and pain threshold were calculated as the averages of the first and second sensory threshold, and of the first and second pain threshold, respectively. The amplitude of the pain stimulus (US+) was set at 1.8 x pain threshold, and the amplitude of the tactile stimulus (US-) was chosen as the middle value between sensory and pain thresholds. For example, if the sensory threshold was 3 mA and the pain threshold was 17 mA, then the amplitude of US+ was 31 mA, and that of US- was 10 mA.

The experiment entailed two phases: an acquisition phase and a test phase (see Figure 1 for graphical representation of the experimental design). During the experiment, subjects were sitting in a comfortable chair with closed eyes. They heard three harmonic tones presented binaurally by means of pneumatic earphones (3M E-A-RTONE). One of them (Standard) consisted of the frequencies 150, 300, 600, 1200, and 2400 Hz. The other two were referred to as Deviant 1 (100, 200, 400, 800, 1600 Hz) and Deviant 2 (250, 500, 1000, 2000, 4000 Hz). The only instruction was to sit still and to listen to the tones.

**Figure 1.**
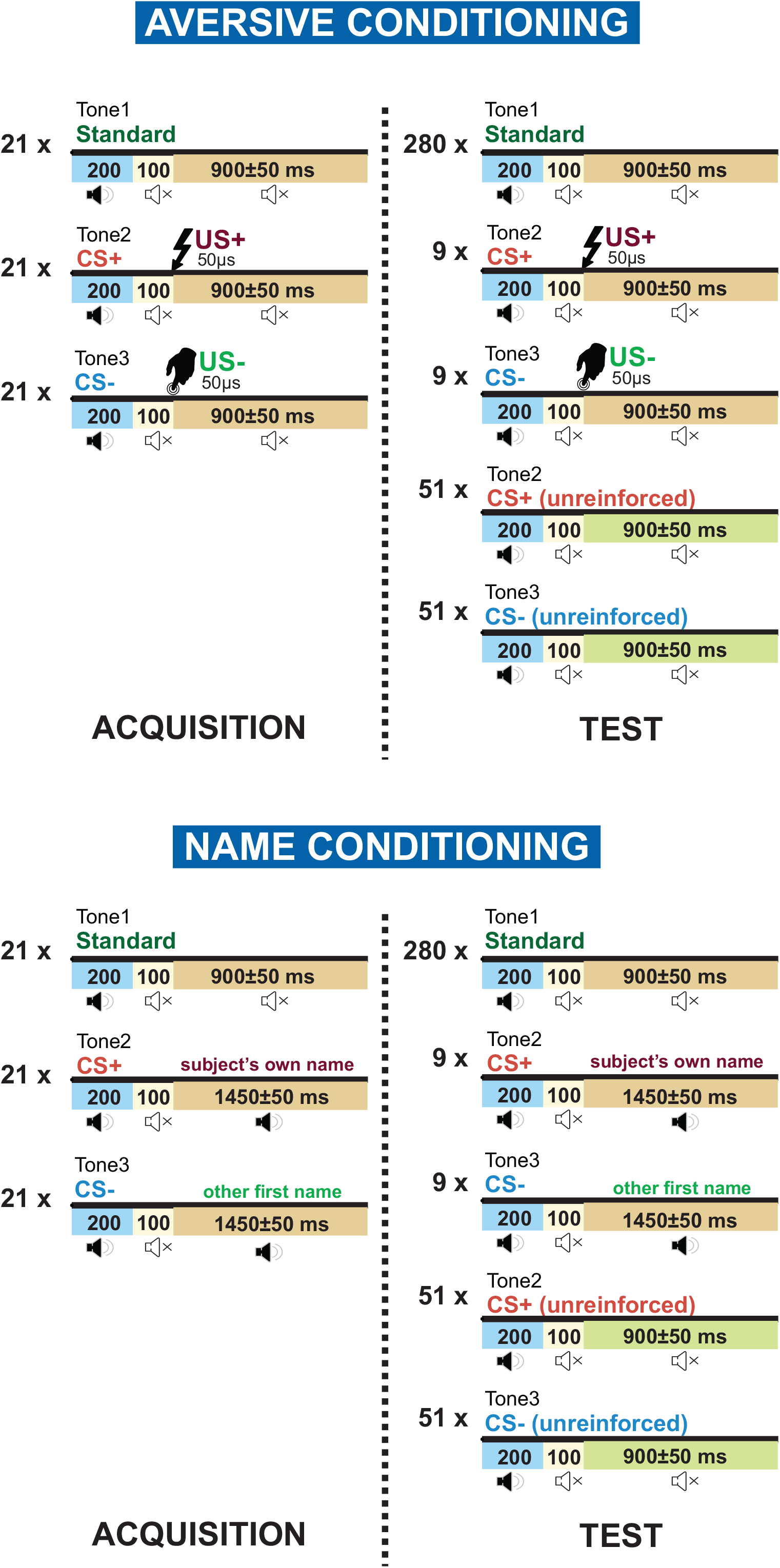
Experimental design. Top panel: Aversive conditioning experiment. In the acquisition phase three types of auditory stimuli were presented: Standard (never being reinforced), CS+ (a tone paired with painful US+), CS-(a third tone paired with a weak electrical shock, US-). In the test phase CS+ and CS-tones were presented each 51 times without reinforcement and 9 times with reinforcement, while Standard was presented 280 times. Unreinforced CSs were included into analyses. Bottom panel: Name conditioning. The design is similar to Aversive conditioning but own names of the participants were used as US+, the other acoustically similar names were used as US-. The average duration of the own name and the other names was 669 ms and 676 ms respectively.

In the acquisition phase the three sounds were presented each 21 times in a random sequence. With 100 % reinforcement rate one of the two Deviants (CS+) was randomly selected to be paired with the pain stimulus (US+), and the other Deviant (CS-) was similarly paired with the tactile stimulus (US-). The details of the pairing are presented in Figure 1. The Standard was never paired with any other stimulus.

The test phase started immediately after the end of the acquisition phase. It was an oddball paradigm where the Standard was presented 280 times, and the Deviants, 60 times each. The order of the presentation was random except that the same Deviant could not be delivered more than two times in a row. Tone duration was 200 ms with stimulus-onset asynchrony (onset-to-onset) varying between 1150 and 1250 ms. Tone intensity was kept about 65 dB SPL.

The test phase followed the procedure of partial reinforcement with 15 % reinforcement rate. Each Deviant was randomly followed by the corresponding electrical stimulus on nine of the 60 presentations, but presented without an electrical stimulus on the remaining 51 trials (Figure 1). Only *the unreinforced trials* were included into analysis.

The average intensity of the pain stimulus (US+) was 39.7±15.9 (range 17-75) mA, and the average intensity of the tactile stimulus (US-) was 13.2±4.4 (range 6-23) mA.

#### Name conditioning

The design of the Name conditioning experiment was identical to Aversive conditioning (see Figure 1) except different US and, accordingly, modified time intervals. In the Acquisition phase a harmonic CS+ tone (300, 600, 120, 2400, and 4800 Hz) was paired with the own name of the corresponding participant (SON), a CS- tone (195, 390, 780, 1560, and 3120 Hz) was randomly paired with three other familiar names (OFN), and Standard (495, 990, 1980, 3960, and 7920 Hz) was presented without any relation to other stimuli. Tone-Name association was counterbalanced between participants. The onset-to-onset interval within a pair tone-name was 300 ms. The onset-to-onset intervals between pairs were 1700-1800 ms, and after standards (which were not accompanied by any word) these intervals were 1150-1250 ms. The average duration of the own name and the other names was 669 ms (SD = 9 ms) and 676 ms (SD = 12 ms) respectively (t = 0.78, p = .44). Other names originated from the same pool of the most frequent German names used for each subject’s own name, and always contained the same number of syllables as the own name. The test phase included partial reinforcement with 15 % reinforcement rate.

### EEG recording

A 64-channels EEG system with active electrodes (ActiCHamp, Brain Products) was used for the recording. The electrodes were placed according to the extended 10-20 system with Cz channel as the online reference and Fpz as the ground electrode. The level of impedance was maintained below 20 kOm. The sampling rate was 1000 Hz.

### ERP analysis

EEGLAB (Delorme & Makeig, 2004) was used for data preprocessing. Each recording was filtered by applying 0.1 Hz high-pass and 45 Hz low-pass filters. Bad channels were replaced by means of spherical interpolation. Data fragments contaminated by high amplitude artefacts (>300 μV) were dismissed. Then, the Independent Component Analysis was performed using the AMICA algorithm (Palmer, Kreutz-Delgado, & Makeig, 2012). Components clearly related to eye movements were removed. Additionally, components that were mapped onto one electrode and could be clearly distinguished from EEG signals were subtracted from the data. After this, data were re-referenced to common reference and epoched in [-200 800 ms] intervals, where [-200 0] interval was used for baseline correction. Epochs still containing artefacts were visually identified and discarded. Finally, before entering the statistical analysis data were re-referenced to average mastoids.

For the analysis of ERP to CS, mean amplitudes of N1, P2, P3a, P3b and LPP were computed in time windows of 70-110, 120-180, 180-250, 290-380 and 400-700 ms respectively. These time windows were chosen on the basis of group average collapsed across all experimental conditions. The data then entered a repeated-measures ANOVA with factors Channel (3 levels: Fz, Cz and Pz) and Condition (2 levels: CS+ and CS-). In order to assess the differences between the Aversive conditioning and Name conditioning experiments we added a factor Experiment (2 levels) and repeated the ANOVA. The analyses did not include ERP to Standard, because its comparison with Deviants simply revealed the well-known ERP oddball effects.

### Time-frequency analysis

Preprocessing steps for time-frequency and connectivity analysis were identical to those in the ERP analysis with two exceptions: 1 Hz high-pass filter was applied, and epochs were defined as [−1500 2500] ms to avoid edge artefacts. All epochs were then converted into current source density (CSD) by means of CSD toolbox (Kayser, 2009). We used spherical spline surface Laplacian (Perrin, Pernier, Bertrand, & Echallier, 1989) with the following parameters: 50 iterations; *m* = 4; smoothing constant λ = 10^−5^ (for detailed description of the procedure see Tenke & Kayser, 2005). This method sharpens EEG topography, diminishes volume conduction effects and has been found to be useful in performing a synchronization analysis (Cavanagh, Frank, Klein, & Allen, 2010; van Driel, Knapen, van Es, & Cohen, 2014). Moreover, the reduction of volume conduction effects by application of CSD transformation may lead to more accurate characterization of functional connectivity (Cavanagh, Cohen, & Allen, 2009; Srinivasan, Winter, Ding, & Nunez, 2007).

The power spectrum of CSD-EEG time series in each epoch was convolved with power spectrum of a set of complex Morlet wavelets and then the inverse fast Fourier transform was taken. The wavelets were defined as: *e*^−*i*2*πtf*^ *e*^−*t*^2^/(2*σ*^2^)^, where *t* is time, *f* is frequency, and *σ* defines the width of each frequency band, set according to *n/(2πf)*, where *n* is the number of wavelet cycles. The frequency *f* increased from 1 to 45 Hz in 45 linearly spaced steps, and the number of cycles *n* increased from 3 to 12 in 45 logarithmically spaced steps. From the resulting complex signal, the power of each frequency at each time point was obtained. The power was baseline-normalized to dB in respect to [−400 −100] ms interval.

To define regions and time-frequency windows of interest we applied a multistep procedure. First, in the Aversive conditioning experiment we compared all reinforced trials (US+ and US-), on the one hand, and a representative group of non-reinforced trials (Standards in the acquisition phase plus non-reinforced trials that preceded reinforced trials in the test phase), on the other hand. We conducted the time-frequency analysis as described above (see Figure 3A) and ran cluster-based permutation tests in the channel-time-frequency space by means of Fieldtrip toolbox (Maris & Oostenveld, 2007; Oostenveld, Fries, Maris, & Schoffelen, 2011) to obtain a reinforced minus non-reinforced difference. As expected, the largest difference was found in the right somatosensory area (channels C2, C4, FC4, CP4) because the reinforcement was always delivered at the left wrist. These channels were characterized by very strong suppression of alpha and beta activity in reinforced trials. Based on these data, two regions of interest (ROI) were defined (marked in Figure 3A): the right somatosensory ROI (C2, C4, FC4, CP4) and the symmetrical left ROI (C1, C3, FC3, CP3). After this we reran the permutation tests using the right somatosensory ROI contrasting US+ vs Standards and US- vs Standards. The intersecting time-frequency window representing features common for both US- and US+ was used to define frequencies of interest.

After the limits of the investigated window of interest in space (i.e., two groups of electrodes over the left and right somatosensory cortex) and frequency (13-19 Hz) were delineated, at the last step the time limits of this window (240-600 ms after CS onset) were defined in the same way (see Figure 3B^2^). The average spectral power in this window entered a repeated-measures ANOVA with within-subject factors Side (left vs right ROI) and Condition (CS+ vs CS-).

### Connectivity analysis

We estimated phase connectivity by means of the debiased weighted phase-lag index (dwPLI; Vinck, Oostenveld, van Wingerden, Battaglia, & Pennartz, 2011). dwPLI is robust to the effects of volume conduction and uncorrelated noise and is not affected by the number of trials in each condition. In order to identify the activity of the auditory cortex, we applied CSD transform to the ERP data. The sources of the N1 components were found at P7/8, T7/8, TP9/10, and P7/8 electrodes (Figure 4A). Because it is known that N1 is originated mainly in the auditory cortex (e.g., Pantev et al., 1995), we assumed that the above electrodes reflect the activity of this cortical area. Therefore, they were used in the connectivity analysis as representing the auditory system. dwPLI was calculated for each possible pair of electrodes between the left somatosensory ROI and the left auditory ROI, the same was done for the right ROIs.

Cluster-based permutation tests were run for an exploratory analysis of the differences in connectivity between CS+ and CS-. First, dwPLI in the left somatosensory-auditory ROI over each frequency and time point entered the test with 5000 permutations. Statistical significance was set at p<0.05, after cluster-based correction. Then the test was repeated for the right ROI.

The time-frequency, connectivity analyses and permutation tests were performed by means of the Fieldtrip toolbox.

## Results

### Event-related potentials

#### Aversive conditioning

As can be seen in Table 1, the amplitudes of *P2* and *P3b* components did not significantly differ between CS+ and CS- (no significant effect of Condition or interaction with Channel). *N1* tended to be more negative in response to CS+ (p = 0.08). The amplitude of N1 was higher at Fz and Cz than at Pz, and the opposite was true for P3b, yielding significant Channel effects (see Figure 2).

**Table 1.**
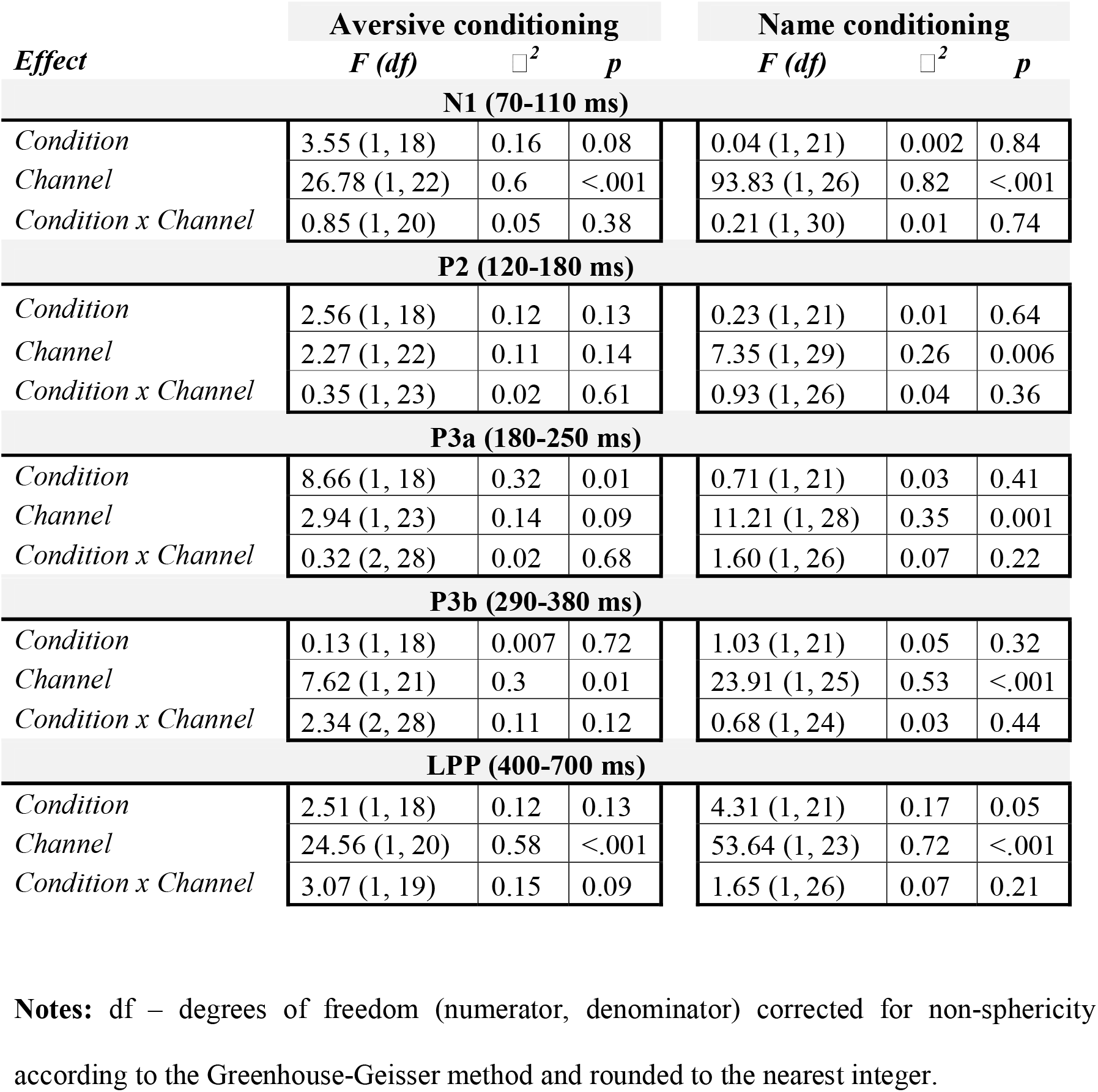
Statistics for ERP analysis

**Figure 2.**
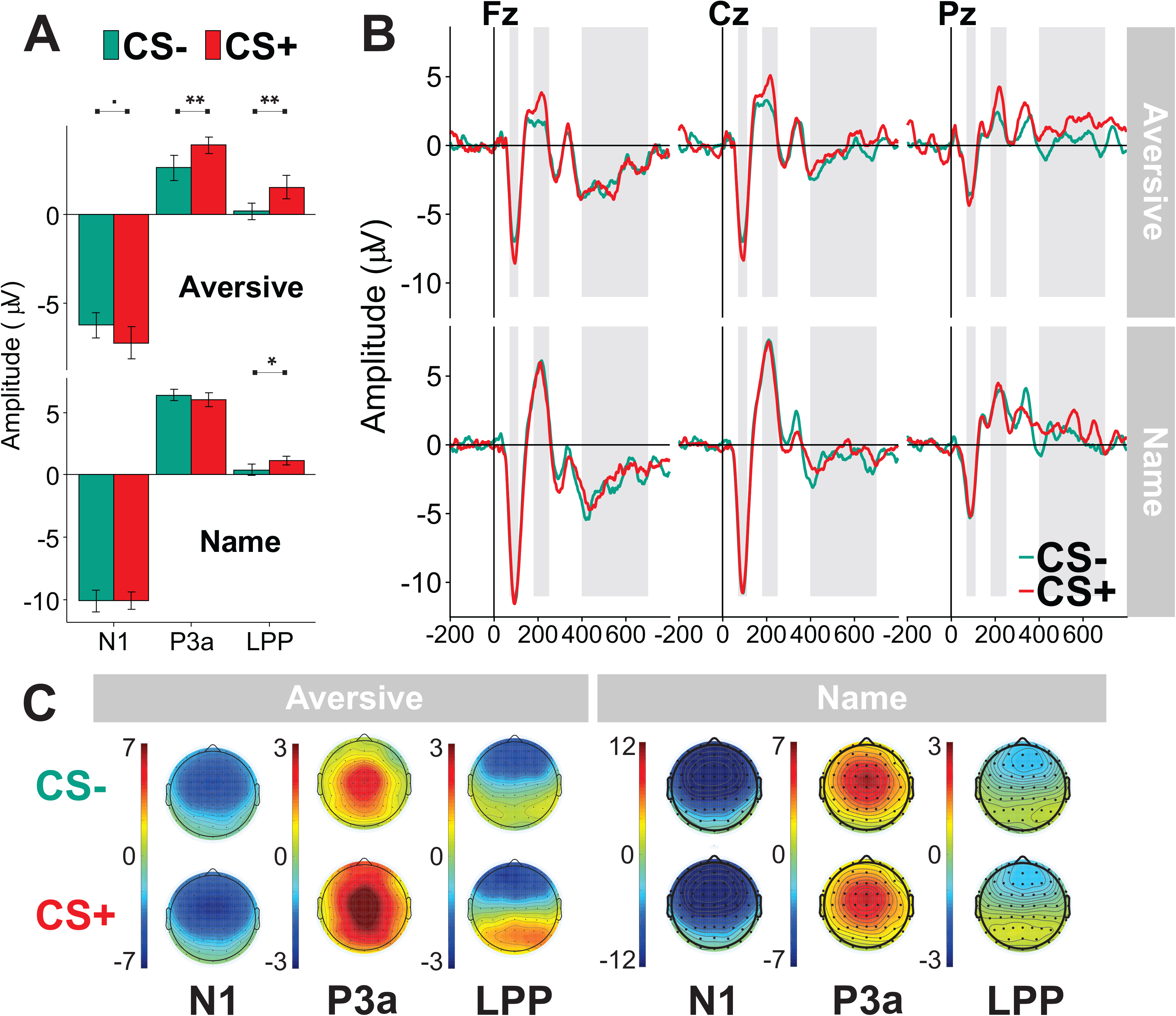
(A) Average amplitudes in the N1 (70-110 ms), P3a (180-250 ms) and LPP (400-700 ms) time windows at Fz, Cz and Pz sites, respectively. Error bars show standard errors of mean. (B) Event-related potentials (referenced to average mastoids) in the CS+ and CS-conditions. Grey areas mark N1, P3a, and LPP time windows (left to right). (C) Corresponding topograms averaged within the components’ windows. · p < 0.1, * p < .05, ** p < .01

After conditioning, *P3a* was larger to CS+ than to CS- (main effect of Condition). As expected, the amplitude of the LPP was larger at Pz than at Cz and Fz (main effect of Channel). Because the ANOVA revealed a tendency to a Condition by Channel interaction, and because a common practice is to analyse the LPP only at Pz (Bacigalupo & Luck, 2018; Liu, Huang, McGinnis, Keil, & Ding, 2012), we conducted an additional ANOVA at the Pz electrode using Condition as a single within-subject factor. The analysis showed a larger LPP amplitude in to CS+ than to CS-(F(1, 18) = 12.31, p = 0.003, □^2^ = .41). Similar analyzes at Fz and Cz did not yield significant effects.

#### Name conditioning

The data are summarized in Table 1 and Figure 2. The only significant effect including the factor Condition was larger LPP amplitude to CS+ than CS-. Like in Aversive conditioning, we also analyzed LPP at Pz alone. The analysis revealed a significantly larger LPP to CS+ than to CS- (F(1, 21) = 5.29, p = 0.03, □^2^ = .20).

An analysis including all subjects who participated in both experiments revealed that LPP effect was not moderated by Experiment (no significant main effect or interaction with Experiment, p > 0.5). A strong main effect of Condition on the LPP at Pz confirmed the results obtained in each experiment (F(1, 16) = 15.72, p = 0.001, □^2^ = .50). The main effect of Experiment was highly significant for the amplitudes of N1 (F(1, 16) = 12.49, p = 0.003, □^2^ = .44) and P3a (F(1, 16) = 53.63, p = <.0001, □^2^ = .77) indicating that these components were larger in Name than in Aversive conditioning. The difference in the P3a amplitude between CS+ and CS-was significant in Aversive conditioning but not in Name conditioning, resulting in a significant Experiment x Condition interaction: F(1, 16) = 4.96, p = 0.04, □^2^ = .24). No significant main effect of Experiment or interaction with Condition was found for P3b.

### Time-frequency analysis

#### Aversive conditioning

We found a significant interaction between Condition and Side (F(1, 18) = 9.71, p = 0.006, □^2^ = .35) in the extracted time-frequency window (13-19 Hz, 240-600 ms). Subsequent ANOVAs for separate ROIs showed stronger lower beta suppression in the CS+ than in CS-condition in the right somatosensory ROI (main effect of Condition: F(1, 18) = 5.46, p = 0.03, □^2^ = .23). No such effect was obtained in the left ROI: F(1, 18) = 0.49, p = 0.49, □^2^ = .03.

#### Name conditioning

In Name conditioning experiment no significant effects were found.

The analysis of the combined data of both experiments yielded a highly significant three-way Experiment x Condition x Side interaction (F(1, 16) = 13.51, p = 0.002, □^2^ = .46) thus confirming that the lateralized beta suppression took place only in the Aversive conditioning experiment (see Figure 3 C,D).

**Figure 3.**
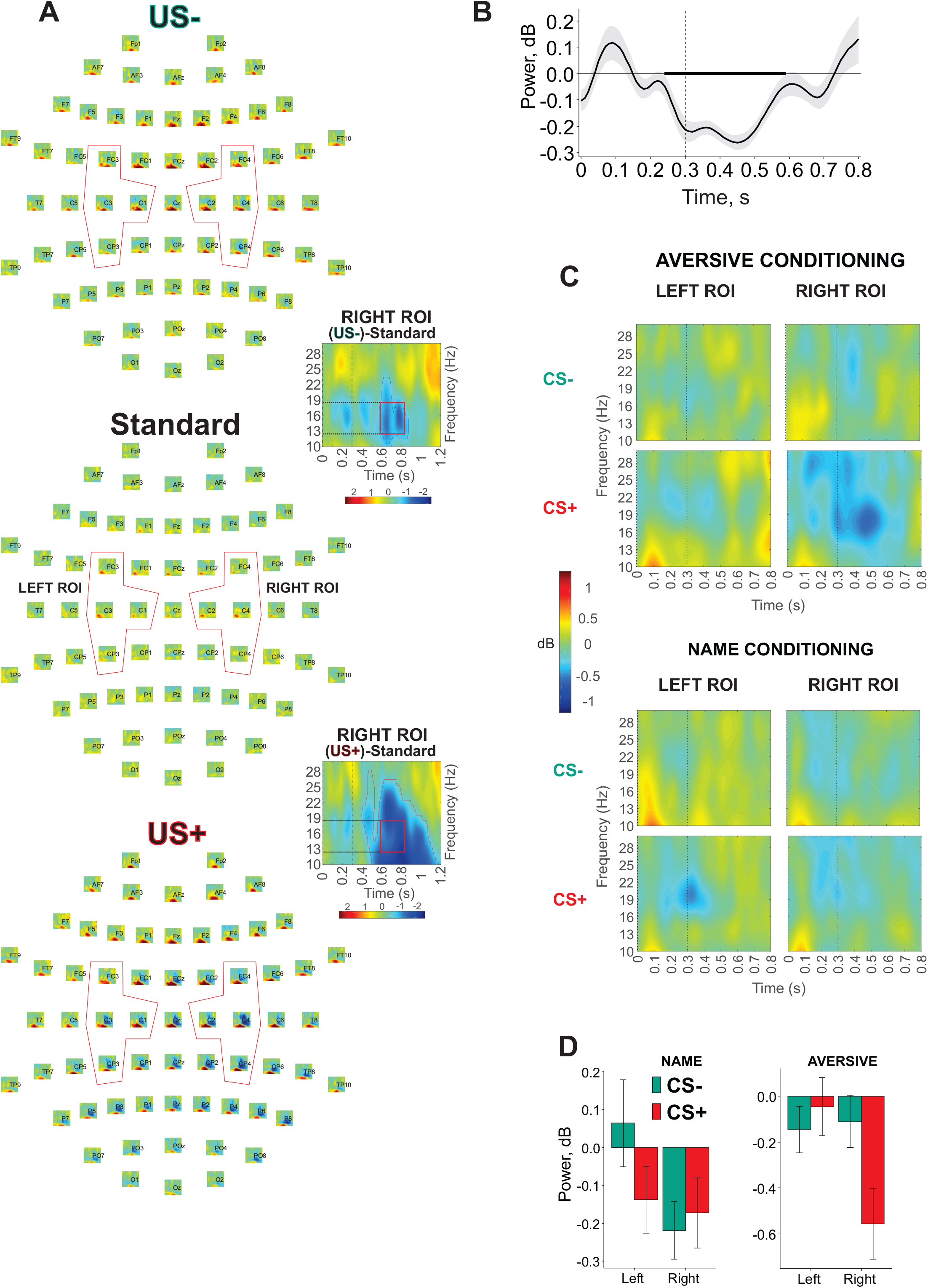
(A) Topographical time-frequency (TF) plots for Standard, US- (weak electrical shock), and US+ (strong painful electrical shock) stimuli. The Left and Right ROIs are marked. Time scale: −100 −1100 ms. Frequency scale: 3-30 Hz. TF plots are based on the results of cluster permutation tests comparing US- and Standard trials, or US+ and Standard trials, in the right ROI. Blue and red sections mark statistically significant clusters in the US- TF plot and US+ TF plot, respectively. Red box in both TF plots shows the window used to define frequencies of interest in the main analysis comparing unreinforced CS+ and CS- trials. (B) Spectral power in time averaged over all ROIs, all conditions in frequency band defined in the previous step (13-19 Hz). Bold line marks the time window of interest. Grey shading is the standard error of mean. (C) TF plots in the Conditions and ROIs (CS-- top row, CS+ -bottom row, Left ROI – left column, Right ROI – right column). Top panel: Aversive conditioning experiment. Bottom panel: Name conditioning experiment. (D) Bar plot of the average spectral power in the Left and Right somatosensory ROIs in the CS+ and CS-conditions during the 240-600 ms time interval. Error bars show the standard errors of mean.

**Figure 4.**
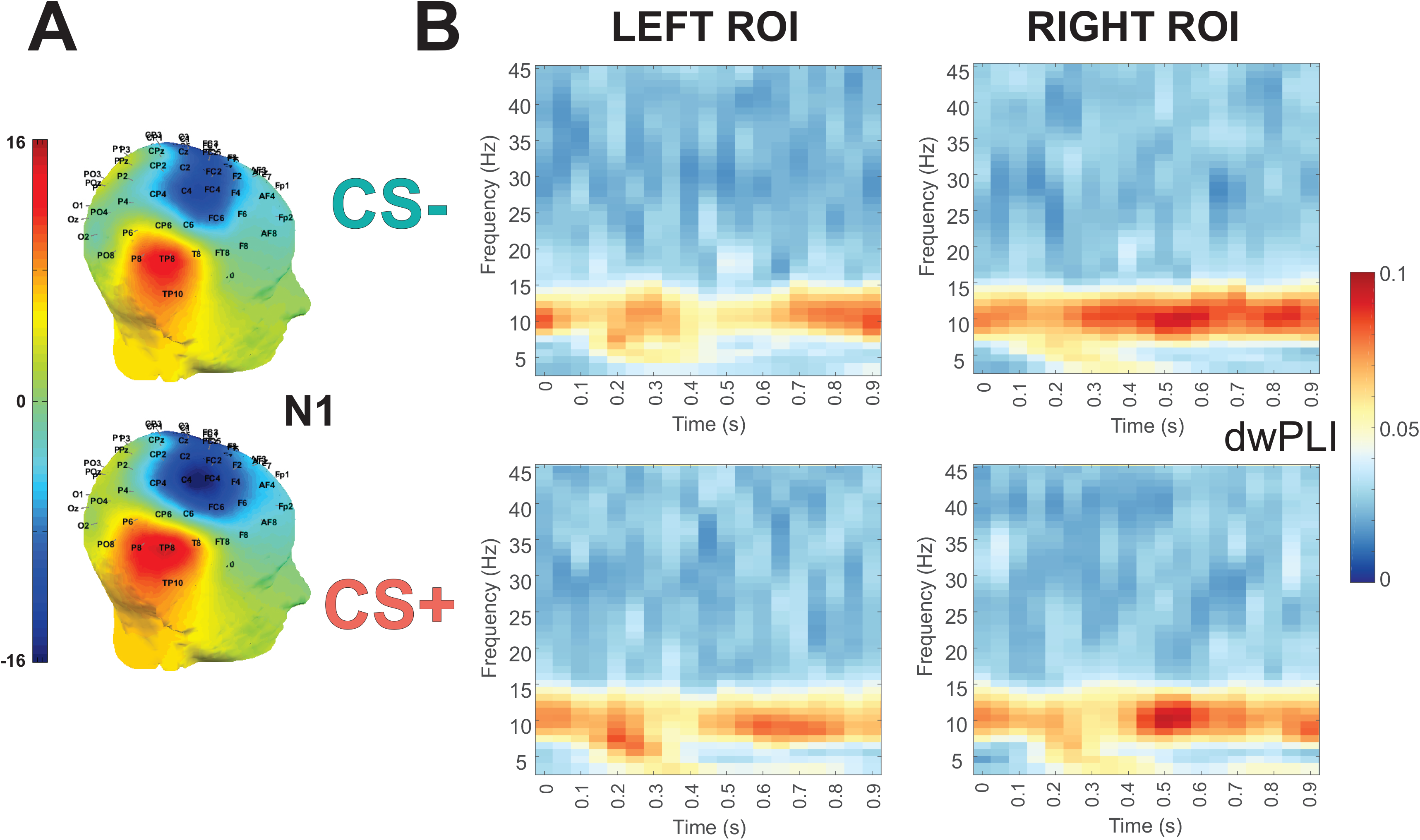
(A) The topographical representation of N1 component of ERP after CSD transformation was used to define the auditory ROI (TP9, TP10, P7, P8, TP7, TP8, T8, T7) (B) Time-frequency plots represent results of the analysis of functional connectivity between the left somatosensory and auditory ROI (left column) and between the right somatosensory and auditory ROI (right column). CS- - top row, CS+ - bottom row. dwPLI was used as a measure of connectivity.

### Connectivity analysis

Two analyses were carried out. First, we performed an exploratory search for any signs of increased connectivity in the CS+ condition within the time-frequency domain of 0-800 ms and 1-45 Hz. No significant clusters of increased connectivity in CS+ were found. Second, we followed Miltner et al. (1999) who obtained a clear effect in the gamma frequency band in the domain of 37-43 Hz. Thus we reran the analysis of dwPLI for this particular frequency range and again did not find any significant effect.

## Discussion

### Event-related potentials

First of all, we explored the Aversive conditioning paradigm as an anchor point to test which conditioning effects can be obtained in this situation with a very salient US. Then we compared it with a possibly weaker (but more suitable for sensible populations) paradigm where own names of the participants were used as US. In the test phase of Aversive conditioning, we found a larger P3a amplitude in response to CS+ than CS-. Unpleasant sounds can capture involuntary attention, thus increasing P3a without affecting earlier components of ERP (Thierry & Roberts, 2007). In our case P3a can be seen as a sign of involuntary attention to meaningful and emotionally laden stimuli associated with electrical shocks.

The amplitude of the late positive potential (LPP) was also larger to CS+ than CS-. The LPP was shown to be an electrophysiological index of emotional processing (Liu et al., 2012). A similar LPP waveform was obtained in an experiment using IAPS pictures as US (Schupp et al., 2000). Previous studies reported an increased LPP in response to emotionally charged auditory stimuli such as emotional prosody, emotional sounds from the International Affective Digitized Sounds database (Hettich et al., 2016; Masuda et al., 2018; Schirmer & Gunter, 2017), words uttered with emotional intonation (Paulmann, Bleichner, & Kotz, 2013), and words with emotional connotation (Hatzidaki, Baus, & Costa, 2015). The enhanced LPP amplitude may reflect cognitive evaluation and categorization of affective stimuli (Ito & Cacioppo, 2000; Olofsson et al., 2008), or their encoding in memory (Olofsson et al., 2008).

The comparison with Name conditioning demonstrated that LPP effect was present in both experiments but P3a effect characterized only Aversive conditioning. Although we found no significant interaction Experiment x Condition, amplitude of N1 tended to be larger to CS+ than CS-in Aversive conditioning but not in Name conditioning. Significant ERP CR can appear as early as 55 ms after CS presentation (Stolarova, Keil, & Moratti, 2006). Aversive conditioning was also found to induce early changes in visual and auditory P1 (Kluge et al., 2011; Muench, Westermann, Pizzagalli, Hofmann, & Mueller, 2016), in visual and auditory N1/P2 components (Bröckelmann et al., 2011; Kluge et al., 2011), as well as in P3 (Baas et al., 2002; Christoffersen et al., 2017; Franken et al., 2011; Wong et al., 2004) and the LPP (Bacigalupo & Luck, 2018; Wong et al., 2004) in visual paradigms. Miskovic & Keil (2012) argued that occurrence of the early effects originating in the primary sensory cortex is related to the number of CS-US pairings. We assume that the stronger the learned association between CS and US, the earlier the effect. Another factor that may affect the latency of the effect is the salience of US. Strong painful US after 21 pairings with CS in the acquisition phase resulted in differential CR in P3a and LPP, and, speculatively, would affect even earlier components with a larger number of pairs. In contrast, a weaker name US in the same condition was only able to modulate the latest stages of stimulus processing, namely the LPP. When, however, the number of acquisition trials is increased, name conditioning paradigm also shows an earlier (P3a) effect (Kotchoubey & Pavlov, 2017).

We further asked whether ERP effects typical for visual CS are generalizable to the auditory modality. Waschulewski-Floruss et al. (1994) found no differential CR in auditory CS – nociceptive US classical conditioning, and Kluge et al. (2011), who used magnetoencephalography, reported changes in P1m, N1m and P2m but no later ERP effects. We successfully reproduced P3 and LPP effects in an auditory CS – pain US conditioning paradigm. This observation suggests that similar mechanisms responsible for generation of late cognitive ERPs in visual and auditory systems.

### Time-frequency analysis

Although auditory stimuli were presented binaurally, they elicited a distinctly lateralized suppression of the lower beta rhythm (13-19 Hz). This lateralization can be regarded as a result of continuous pairing between auditory (tones) and somatosensory (lateralized electrical shock) stimuli in the preceding acquisition phase.

Outside the framework of conditioning, the suppression of alpha and beta oscillations in anticipation of tactile events (e.g. van Ede et al., 2014) was interpreted as an increase of neuronal excitability in the somatosensory cortex aiming at improving task performance (van Ede, de Lange, & Maris, 2012). Since in our experiment no task was given, a different interpretation is required. It was proposed that the observed effect may represent a preparatory mechanism serving to adjust the somatosensory system, thereby reducing the noxious impact of the electrical shock (Miskovic & Keil, 2012; Wik, Elbert, Fredrikson, Hoke, & Ross, 1996).

On the other hand, a later portion of the observed local beta suppression might reflect, not only the prediction of the pain stimulus, but also a response to prediction error after this stimulus has not been presented. Such effect of the omission of anticipated somatosensory stimuli on cortical oscillations was demonstrated in a recent MEG study (Andersen & Lundqvist, 2019). Another MEG experiment revealed conditioning-induced suppression of beta activity in the contralateral somatosensory cortex to CS+ alone (Moses et al., 2010). The peak latency of the effect in that study was between 150 and 300 ms after the omission of the anticipated US. In the current study the peak latency was between 100 and 250 ms after US omission (Figure 3). However, our data provide no evidence that the suppression of beta, starting 60 ms before expected US, substantially changes after this point; therefore, the data do not permit a clear dissociation between the processes of expectation and its violation. Possibly, experiments with longer CS-US intervals may yield clearer results in this respect.

### Connectivity

In a fMRI experiment Greening, Lee, & Mather (2016) used auditory CS and demonstrated activation of the primary somatosensory cortex in trials in which aversive electrical shock (US) was omitted. The activation was accompanied by increased functional correlation between the somatosensory cortex and amygdala and between the somatosensory and the auditory cortex. This finding may indicate a mechanism of multisensory integration underlying associative learning. In search of EEG manifestations of this mechanism we applied a connectivity analysis testing for increased phase synchronization between the auditory (representing CS) and the somatosensory (US) cortex. The initial exploratory analysis across all time points and frequencies resulted (after a correction for multiple comparisons) in a zero finding. Then, in search for a possible hypothesis, we followed Miltner et al. (1999) who found fear-conditioning induced coherence between somatosensory and visual cortical areas in a specific range of 37-43 Hz. Our results did not replicate this finding. In the light of the recent data showing no reliable relationship between local field potentials and scalp EEG coherence (Snyder, Issar, & Smith, 2018), this is not surprising. An important difference between our experiment and that of Miltner et al. (1999) is a much larger CS-US interval in the latter. On the other hand, Miltner et al. (1999) did not provide a convincing justification of the selected time-frequency window where the effect was found. More research is needed to validate the use of EEG phase synchronization to reveal possible mechanisms of associative learning.

### Limitations

First, our study used a limited number of CS-US acquisition trials. More pairings might strengthen the conditioning effects, particularly the weaker effects in the Name conditioning experiment. Second, as we tried to make brief paradigms appropriate for individuals with limited cognitive abilities, we used rather short CS-US intervals. Also this parameter may have considerable effects on EEG measures. Finally, we did not explore extinction of conditioned responses that would take place when partial reinforcement is switched off. Because the necessity of averaging, an analysis of extinction using ERPs is a specific methodological task that should be dealt with in a separate study.

## Conclusions

We tested a novel experimental design to study associative learning where classical conditioning was combined with passive oddball. The paradigm does not demand any instruction or training of participants and lasts for 10 min only. We showed that aversive conditioning in this paradigm strongly influences brain activity; therefore, the learning process can be detected by the EEG even in the absence of any behavioral index. Aversive conditioning using electrical shocks resulted in a local suppression of lower beta activity over the hemisphere contralateral to expected electrical shocks. The increase of the amplitudes of P3a and the LPP to conditioned stimuli can represent signatures of enhanced attentional and emotional significance of these stimuli. These LPP and P3a effects, earlier found only in visual paradigms, generalize to the auditory sensory system.

## Acknowledgement

The study was supported by the German Research Society (Deutsche Forschungsgemeinschaft), Grant KO-1753/13.

## Footnotes

1 In the literature there is a tendency to use the terms „aversive conditioning” and “fear conditioning” as synonyms. We prefer to speak about “fear conditioning” only in those cases, in which independent data indicate that subjects really experienced fear. In contrast, a conditioning procedure using aversive (potentially fear-generating) stimuli can be referred to as “aversive conditioning” regardless of which kind of emotion (fear, anxiety, disgust, etc.) was experienced, and in what extent.

2 We also applied other criteria to define the time window. All of them yielded the same result.

