## Supplementary for "Classical conditioning in oddball paradigm: A comparison between aversive and name conditioning"

**Supplementary materials**

***Methods***

In order to investigate correlations between LPP and personality we used State Trait Anxiety Inventory (STAI)-trait, the NEO Five-Factor Inventory (NEO-FFI) at the beginning of the experimental session and STAI-state before and after the experiment. We calculated Spearman rank-order correlations by means of Hmisc R package (Harrell Jr & Dupont, 2008).

### In Aversive conditioning experiment one subject did not complete STAI-trait questionnaire and two did not complete NEO-FFI. In Name conditioning one subject did not complete STAI-trait questionnaire and NEO-FFI. They were excluded from the correlational analysis.

***Personality***

**Aversive conditioning**. The amplitude of the LPP in the CS+ condition was negatively related to STAI-trait (rho = -0.58, p = 0.009) and neuroticism as a Big Five trait (rho = -0.67, p = 0.002, see Figure S1 A,B). The correlation between STAI-trait and Neuroticism was rho = 0.8, p = 0.0009. No significant correlations were found between the LPP and other subscales of NEO-FFI or amongst the subscales. The lack of correlations between the subscales can be interpreted as a measure of the reliability. The difference between STAI-state before and after the conditioning procedure did not significantly correlate with the LPP.

**Name conditioning.** No significant correlations between CS+ LPP and the personality traits were found (see Figure S1 C,D).

As expected, all correlations of the LPP with non-emotional personality factors of the Big Five model (i.e., Extraversion, Openness, Conscientiousness, and Agreeableness), were around zero.

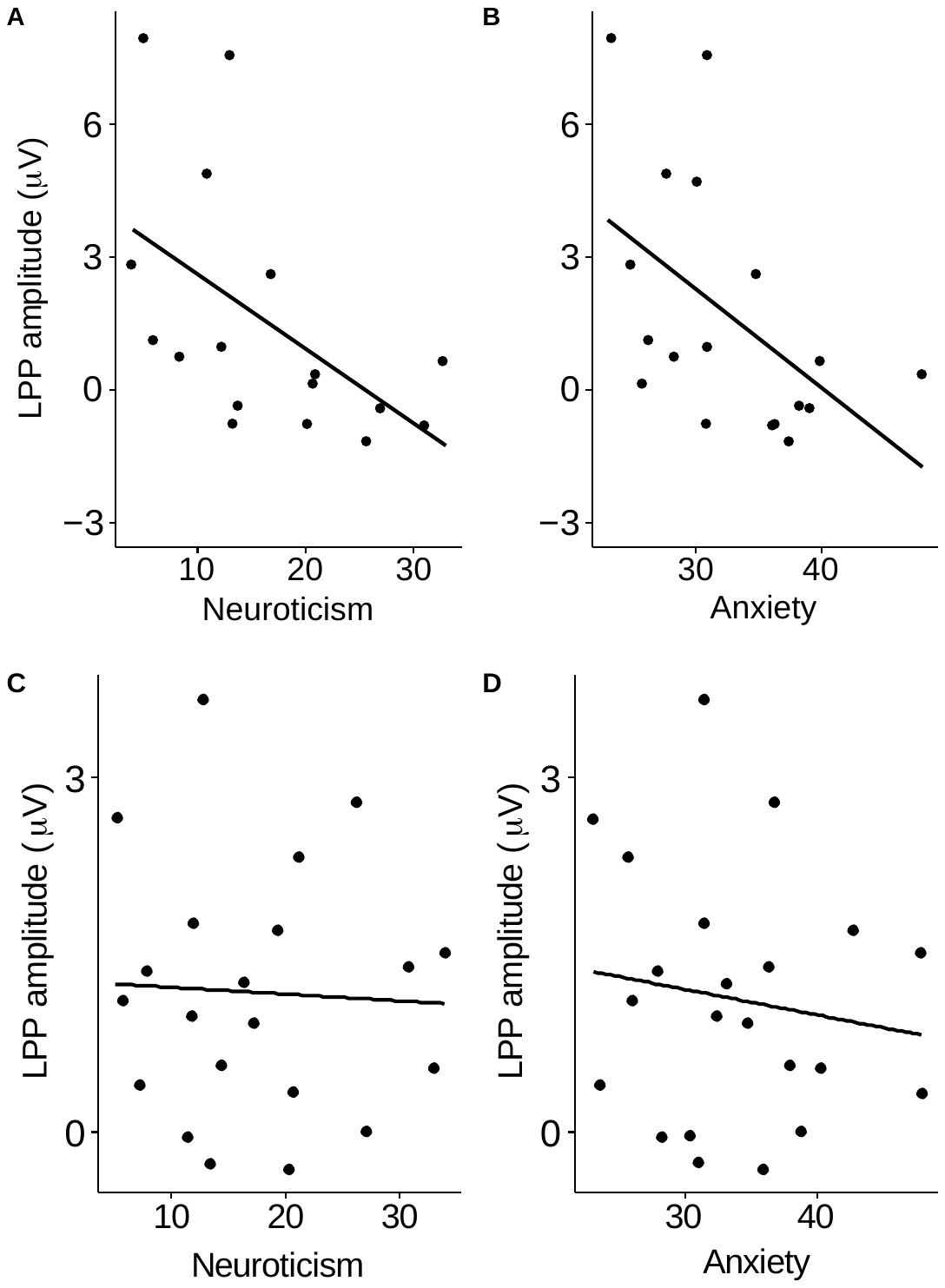

Figure S1 – Correlations between personality traits and the amplitude of the late positive potential. A: Correlation of LPP amplitude and neuroticism in Aversive conditioning. B: Correlation of LPP amplitude and trait anxiety in Aversive conditioning. C and D: the same in Name conditioning.

***Discussion***

“[A]ffective ERP waveform variability across individuals has received very little consideration” (Olofsson et al., 2008, p. 12). In the current study the amplitude of the LPP was inversely related to individual traits that reflect emotional aspects of personality. We should stress that the sample size in our study is relatively low for studies on personality; therefore the results should be treated as preliminary. Nevertheless, a similar negative relationship between anxiety and the amplitude of the LPP was found by Holmes, Nielsen, & Green (2008) who observed attenuated LPP in response to fearful facial expressions in high-anxiety as compared with low-anxiety individuals. However, in another study neuroticism and LPP in response to highly arousing unpleasant pictures were positively correlated (Brown, Goodman, & Inzlicht, 2013). Some studies found no correlation between anxiety and LPP (e.g., Taake, Jaspers-Fayer, & Liotti, 2009; Malak, Crowley, Mayes, & Rutherford, 2015).

Olofsson et al. (2008) in their analytic review depict the memory hypothesis as the prevalent interpretation of LPP. According to this interpretation, which capitalizes on the finding of the strong relationship between late ERP components and memory processes (e.g. Azizian & Polich, 2007; Paller, McCarthy, & Wood, 1988), highly arousing affectively negative stimuli immediately activate the amygdala, and this activation leads (among other consequences) to a rapid allocation of cognitive resources for saving the negative event in memory (LeDoux, 2000). If this hypothesis is correct, it might explain that very different stimuli having the only common feature of being emotionally negative, can result in different correlations between LPP and personality. Highly anxious persons (as compared with emotionally stable persons) can better record and save some emotional stimuli but use the opposite strategy (avoidance of memory recording) in respect to other emotional stimuli. This post hoc explanation remains, of course, highly speculative and should be tested in a separate study using, on the one hand, a broad range of negative stimuli, on the other hand, a clinical population of individuals with high levels of anxiety and neuroticism.

***References***

Azizian, A., & Polich, J. (2007). Evidence for Attentional Gradient in the Serial Position Memory Curve from Event-related Potentials. *Journal of Cognitive Neuroscience*, *19*(12), 2071–2081. https://doi.org/10.1162/jocn.2007.19.12.2071

Brown, K. W., Goodman, R. J., & Inzlicht, M. (2013). Dispositional mindfulness and the attenuation of neural responses to emotional stimuli. *Social Cognitive and Affective Neuroscience*, *8*(1), 93–99. https://doi.org/10.1093/scan/nss004

Harrell Jr, F. E., & Dupont, C. (2008). Hmisc: harrell miscellaneous. *R Package Version*, *3*(2).

Holmes, A., Nielsen, M. K., & Green, S. (2008). Effects of anxiety on the processing of fearful and happy faces: An event-related potential study. *Biological Psychology*, *77*(2), 159–173. https://doi.org/10.1016/j.biopsycho.2007.10.003

LeDoux, J. E. (2000). Emotion circuits in the brain. *Annual Review of Neuroscience*, *23*(1), 155–184.

Malak, S. M., Crowley, M. J., Mayes, L. C., & Rutherford, H. J. V. (2015). Maternal anxiety and neural responses to infant faces. *Journal of Affective Disorders*, *172*, 324–330. https://doi.org/10.1016/j.jad.2014.10.013

Paller, K. A., McCarthy, G., & Wood, C. C. (1988). ERPs predictive of subsequent recall and recognition performance. *Biological Psychology*, *26*(1), 269–276. https://doi.org/10.1016/0301-0511(88)90023-3

Taake, I., Jaspers-Fayer, F., & Liotti, M. (2009). Early frontal responses elicited by physical threat words in an emotional Stroop task: Modulation by anxiety sensitivity. *Biological Psychology*, *81*(1), 48–57. https://doi.org/10.1016/j.biopsycho.2009.01.006
